# A Novel Early Onset Spinocerebellar Ataxia 13 BAC Mouse Model with Cerebellar Hypoplasia, Tremor, and Ataxic Gait

**DOI:** 10.1101/2024.10.28.620097

**Authors:** Junxiang Yin, Jerelyn A. Nick, Swati Khare, Heidi E. Kloefkorn, Ming Gao, Michael Wu, Jennifer White, James L. Resnick, Kyle D. Allen, Harry S. Nick, Michael F. Waters

## Abstract

Spinocerebellar ataxia 13 (SCA13) is an autosomal dominant neurological disorder caused by mutations in *KCNC3*. Our previous studies revealed that *KCNC3* mutation R423H results in an early-onset form of SCA13. Previous biological models of SCA13 include zebrafish and Drosophila but no mammalian systems. More recently, mouse models with *KCNC3* mutations presented behavioral abnormalities but without obvious pathological changes in the cerebellum, a hallmark of patients with SCA13. Here, we present a novel transgenic mouse model by bacterial artificial chromosome (BAC) recombineering to express the full-length mouse Kcnc3 expressing the R424H mutation. This BAC-R424H mice exhibited behavioral and pathological changes mimicking the clinical phenotype of the disease. The BAC-R424H mice (homologous to R423H in human) developed early onset clinical symptoms with aberrant gait, tremor, and cerebellar hypoplasia/atrophy. Histopathological analysis of the cerebellum in BAC-R424H mice showed progressive Purkinje cell loss and thinning of the molecular cell layer. Additionally, Purkinje cells of BAC-R424H mice showed significantly lower spontaneous firing frequency with a corresponding increase in inter-spike interval compared to that of wild-type mice. Our SCA13 transgenic mice recapitulate both neuropathological and behavioral changes manifested in human SCA13 R423H patients and provide an advantageous approach to understanding the role of voltage-gated potassium channel in cerebellar morphogenesis and function. This mammalian *in vivo* model will lead to further understanding of the R423H allelic form of SCA13 from the molecular to the behavioral level and serve as a platform for testing potential therapeutic compounds.

## Introduction

Spinocerebellar ataxia (SCA13) is a progressive neurodevelopmental/neurodegenerative disease with major impacts on the daily life of patients, their families, and the health care system. There is no cure for the inherited cerebellar ataxias nor are there effective treatments to slow the progression of its symptoms, cerebellar atrophy, or neuronal degeneration. SCA13 is an autosomal dominantly inherited progressive disorder, the clinical hallmarks of which include ataxic gait, tremor, and cerebellar atrophy ^1–5^. The age of onset, severity, and progression of clinical symptoms in patients with SCA13 are highly heterogenous and attributable to multiple allelic isoforms mutations of *KCNC3* ^1, 6–10^. Phenotypic consistency is seen within allelic groups. In 2006, we reported the link between human mutations in *KCNC3*, encoding the voltage gated Kv3.3 potassium channel, and SCA13. Two mutations were described: R420H, which leads to adult-onset progressive ataxia, and F448L, which results in childhood onset ataxia with cognitive delay^1^. KCNC3 is widely expressed in the nervous system and is especially prominent in Purkinje cells of cerebellar cortex and hippocampal dentate granule cells where they play a critical role in firing high-frequency action potentials ^11, 12^. We previously found that patients with SCA13 harboring the R423H mutation in *KCNC3* show an early-onset, non-progressive form of the disease with delayed achievement of motor and cognitive milestones, dysarthria, tremor and marked gait and/or appendicular ataxia ^10^. In contrast, patients with SCA13 carrying a R420H mutation displayed late-onset progressive cerebellar atrophy; however, genetic modeling of SCA13 by expressing this pathogenic mutant has not been achieved in mammalian model ^9, 13, 14^. *KCNC3* mutation, F448L, leads to an early-onset form of SCA13, with normal mutant protein trafficking to the membrane in cell culture. However, the F448L mutation shows abnormal shifts in activation voltages, further accentuating the phenotypic and pathophysiologic heterogeneity of SCA13 ^9^. Previous mice models with a *Kcnc3* mutation only exhibited very mild phenotypes and there was lack of obvious pathological change in the cerebellum of brain anatomy ^15, 16^. To generate a transgenic SCA13 mouse model and to study the impact of expression of the human *KCNC3* R423H mutant in an *in vivo* system, we used the homologous mouse R424H mutation to mimic the human pathological R423H mutation. We created the transgenic mice by bacterial artificial chromosome (BAC) recombineering to express the full-length mouse *Kcnc3* expressing the R424H mutation (herein referred to as BAC-R424H mice). Our mouse model showed progressive Purkinje cell neurodegeneration, marked cerebellar hypoplasia/atrophy, and motor deficits including tremor, loss of coordination, and gait ataxia. These behavioral and neuropathological changes were observed as clinical characteristics in patients with SCA13 R423H. Thus, this novel murine model will provide an important tool for studying pathogenetic neurodegenerative mechanisms and a platform to test potential therapeutic compounds.

### Materials and Methods

#### Creation of BAC-R424H mice

We obtained a BAC clone (RP23-34H20) from the Children’s Hospital Oakland Research Institute which contains the *Kcnc3* locus. The BAC clone was grown in DH10B, isolated by an alkaline-lysis mini-prep procedure and verified to contain the mouse *Kcnc3* gene using primers designed for four of the *Kcnc3* exons. The BAC DNA was then transferred to the recombineering *E. coli* strain EL250 using electroporation and chloramphenicol selection. BAC integrity was verified by restriction enzymes.

The BAC clone contains the following genes in this order: Napsa, Kcnc3, Myh14, Izumo2, Zfp473 and Vrk3. Primers were designed with relevant restriction sites for PCR of an intron1/exon2 fragment from the BAC clone. This fragment contains 628 bp of intron1 along with a single restriction site for insertion of an FRT-NEO-FRT cassette as well as 752 bp of exon2. The portion containing exon 2 includes the site for insertion of the R424H mutation as well as the silent mutagenesis of an adjacent codon (L423L) for the purposes of generating a diagnostic restriction site. The intron1/exon2 fragment was inserted into a Bluescript vector and mutagenesis was performed using a QuikChange protocol (New England Biolabs). The resulting mutagenesis yielded R424H and a HpyCH4V site at L423L by converting a CTT to CTG, thus maintaining the Leucine at position 423. We utilized standard T7 and T3 primers to PCR from vector ends to confirm the presence of the mutation by DNA sequencing.

For selection purposes, we cloned a ∼1600 bp cassette containing the Neomycin resistance gene flanked by the FRT recombination sites into a HindIII site in intron1 of the Bluescript plasmid. The intron1/exon2 fragment containing the Neo cassette and R424H/L423L mutation was excised from Bluescript using XhoI and NotI. This fragment was then electroporated into the KCNC3 BAC/EL250 bacteria with Kanamycin/Chloramphenicol selection. Temperature dependent induction of antibiotic selected EL250 colonies allows for homologous recombination between the intron1/exon2 mutated fragment and the corresponding region on the KCNC3 BAC. The resulting colonies were then treated with arabinose to activate the FLpase encoded in EL250 leading to excision of the Neo cassette. The presence of the mutation was confirmed by restriction digestion (HpyCH4V) and DNA sequencing. A glycerol stock of bacteria was then sent to the Transgenic Animal Model Core at the University of Michigan. This engineered R424H construct was then microinjected in fertilized mouse oocytes at the University of Michigan Biomedical Research Core. This produced five founder lines (lines A, B, C, D and E). All five of the founder lines demonstrated indistinguishable phenotypes of varying severity providing confidence that insertion site gene disruption nor regional expression regulation was responsible for the observed phenotype. Amongst the five founder lines, line E was difficult to breed, and line C was infertile. Hence these two lines were eliminated from further studies. Lines A, B and D, had indistinguishable phenotypes, based on our observations. Line B showed a milder phenotype but the BAC insertion for this line was on the Y chromosome since all affected mice were males. Line D was the most severely affected line also corresponded with higher BAC copy number, and unfortunately was lost due to reduced fecundity secondary to physical impairments. Line A was utilized for all further experimentation, based on reproducible phenotype in both affected males and females. For simplicity, this founder line A was denoted as BAC-R424H.

FVB/NJ mouse from Jackson laboratory is used for mouse genetic background and transgenic injection. All mice were housed in a temperature and humidity-controlled vivarium, kept on a 12-hour dark/light cycle, and had free access to food and water. All experimental procedures were approved by the Institutional Animal Care and Use Committee of the Barrow Neurological Institute and performed according to the Revised Guide for the Care and Use of Laboratory Animals. All principles of laboratory animal care (NIH publication No. 86-23, revised 1985) were followed.

### Genotyping of BAC-R424H mice

DNA was extracted from a piece of the mouse tail, approximately 0.6 cm in length, through a high salt extraction protocol (40). Briefly, each tail was digested in 600uL of TNES buffer (50mM Tris base, 400mM NaCl, 100mM EDTA, 0.5% SDS) and 18uL of 20mg/mL Proteinase K overnight at 55°C. Once digested,167uL of super saturated 6M NaCl was added to the tubes. The tubes were mixed and centrifuged at 14000 rpm for 5 min, and the supernatant was removed. 600uL of cold 100% ethanol was then added to each tube, mixed well, and centrifuged at 14000 rpm for 2 min. The supernatant was discarded and 1mL of cold 70% ethanol was added, and centrifuged at 14,000 rpm for 2 min. The supernatant was again discarded, and tubes were air-dried, after which the DNA was resuspended in ddH20 until for the PCR reactions. For each reaction 10 μL of Platinum II Green Hs PCR Master Mix (2x) (Invitrogen, # 14001012), 1 μL forward primer, 1 μL reverse primer, and 7 μL nuclease free water were used. These amounts were multiplied by the number of samples and aliquoted into separate tubes at which time 1 μL of the sample supernatant from each mouse was added to its corresponding tube for a total volume of 20 μL. Specific primers (F – 5’ AGA ACC ATA ACC CGT CTC CA 3’, R –5’ CTA TGC CCC CAG ACT CAT TC 3’) designed to differentiate the wild type from the transgenic mice were then used in PCR to identify each genotype, the primers were obtained from Integrated DNA Technologies. The mutation is on the FVB JR1800 genetic background and thus JR1800 was used as our wild type of control. PCR was run using the BIO RAD CFX Connect Real-Time System. The samples were heated to 94 °C for 2 minutes, during step two they were heated to 94 °C for an additional 15 seconds, step 3 heated to 60 °C for 15 seconds, step 4 heated to 68 °C for 15 seconds. Steps 2-4 were repeated 35 times after which they were held at 4 °C until run on a gel. The gel was prepared by adding 2% molecular biology grade agarose into 80 mL of 1x TAE buffer prepared with deionized water. This mixture was heated until the agarose dissolved and then covered and left to cool for 15 minutes. After the cooling period 4 μL of Ethidium bromide was added and then poured into a mold to cool for approximately 30 minutes. 4 μL of GeneRuler DNA Ladder Mix ready-to-use (Thermo Fisher Scientific, # SM0334) was added to the first well and then 6 μL of each sample was added to the subsequent wells. The gel was run in 1x TAE buffer for approximately 30 minutes at 120 V. The gel was then imaged using a LI-COR Odyssey Fc Imager. The R424H mice had a wild type of band at 497 bp and a carrier band at 631 bp.

### Quantitative PCR

Samples preparation and handling follow MIQE guidelines for RT-qPCR^17^. Following brief anesthesia, mice cerebella were quickly harvested and sectioned into 2 mm thick coronal slices using a rodent brain matrices slicer (ASI Instruments, Warren, MI, USA) and then flash frozen in liquid N2. Total RNA was isolated from the slices as individual and homogenized with Trizol reagent (Invitrogen, Cat # 15596026) and purified with RNeasy mini kit (Qiagen, Cat # 74104) following the manufacturer’s instructions within a few weeks of storage. RNA concentrations were determined using Thermo Scientific Nanodrop 1000 Spectrophotometer and integrity was verified by visualization of ribosomal RNA by gel electrophoresis and ethidium bromide staining. To eliminate the potential for genomic DNA contamination in the RT qPCR experiment, all RNA samples are subjected to the amplification protocol prior to reverse transcription with oligo-DdT priming for first strand cDNA synthesis. If an amplicon is generated, the sample is considered potentially contaminated with genomic DNA and is no longer included in any further experiments. Primers were designed with Net primer and Primer Blast software to span exon junctions, with an optimal Tm of 60 1C, and no DNA contamination was detected based on no RT controls. Specific primers used are listed; Calbindin - *F* 5’ GAG TTA TAT GAT CAG GAT GGC AAC G 3’, *R* 5’ GTT CGG TAC AGC TTC CCT CC 3’, Kcnc3 – *F* 5’ TGC CAG GTA TGT GGC CTT CGC 3’, *R* 5’ AAG AAG GGT TCC GTC TCC ACC. cDNA was produced with the SuperScript First-Strand Synthesis Kit (Invitrogen, # 11904018) using oligo dT priming and subsequently utilized for RT-qPCR with SYBR Green master mix (Applied Biosystems, #4309155). 264 ng of total RNA was used for each 20 μl cDNA reaction which was then diluted to 200 μl and 2 μl of diluted cDNA employed for each 25 μl RT-qPCR reaction containing 600 nM of each primer pair. Individual RT-qPCR reactions were carried out in triplicate in an ABI PRISM 7000 Sequence Detection System and relative fold changes were determined using the ΔΔCT method normalized to the reference gene Ppia (cyclophilin A), which also served as the inter-run calibrator. The use of a single reference gene based on the suggestions in MIQE guidelines, demonstrates that the Cq values for Ppia (cyclophilin A) have a standard deviation of 0.36 over the 36 cerebellum tissue samples analyzed as part of this study. This provides the validity for utilizing Ppia as a single reference gene in the analysis of our RT-qPCR data. Briefly, the difference between the quantification cycle (Cq) values of the target and control genes represents the ΔCT value. The difference between the ΔCT value for any given R424h sample and the average of the corresponding wild-type samples generates the ΔΔCT value. 2ΔΔCT provides the relative fold change for each sample compared to wild type, which is normalized to 1.

### Coordination testing

Coordination data was obtained using a Rotarod (IITC Life Science Rotarod Series 8) for mice at age of 3, 6 or 9 months. Mice were acclimatized to the procedure room for 1 h before testing and trained for three consecutive days. Data from four trials per animal were acquired on the fourth day of testing. The test length was set to 10 minutes, and the rod was accelerated from 4 rpm to 40 rpm over 5 minutes. Latency to fall was recorded, and mice were given 15 min of rest between each trial.

### Magnetic Resonance Imaging (MRI) and brain volume measurement

Mice at age of 1, 3, and 8-month-old were scanned under Magnetic Resonance Imaging (MRI) for brain volume measurement. MR images were acquired on a Bruker Biospec 7.0T small animal MR scanner (Bruker Medizintechnik) with a 72mm transmit coil and a surface receive coil. We employed a 3D RARE imaging sequence with RARE factor of 8, 4 averages, yielding 100 μm isotropic voxels with 44 min scan time. A sagittal multiple slice 2D T2-weighted RARE sequence was acquired as a reference image (with RARE factor of 8, 10 averages, TR/TEeff= 5000 ms/60 ms; matrix size of 220 × 100; FOV 22 mm× 10 mm; 25 slices) to visualize sagittal morphology. A gas mixture of oxygen and isoflurane (1–2%) was used for anesthesia maintenance. The respiratory rate and body temperature were monitored constantly with an MR-compatible small animal monitoring system (PC_SAM, model #1025, SA Instruments). The body temperature of the mouse was kept at 36.5°C and the respiratory rate was maintained around 50–65 bpm by adjusting the concentration of the isoflurane gas mixture.

3D MR Bruker files were converted to a NIfTI format in ImageJ v.1.51j8. The converted 3D MR images were skull-stripped using the Brain Surface Extractor (BSE) within BrainSuite v.17a to remove all non-brain material ^18^. Inaccuracies were corrected in all three axes by manually editing the masks using Brain Suite v.17a ^18^. After skull stripping, the cerebella were manually extracted from the brains using a routine allowing manual operator contouring. The volumes of the whole brains and cerebellums were calculated in Brain Suite v.17a using the ROI volume estimation function.

### Tremor analysis

1, 3, and 8-month-old mice were held in a scruff and handled by the same person each time. High speed video recordings (2 trials, 8 seconds per trials, n=5 per group) were taken under ventral view at the frame rate of 240 frames per second, using a Sony Cyber-shot DSC-RX100 IV digital camera. DLT data viewer (https://biomech.web.unc.edu/dltdv) was utilized by a blinded observer for manually marking the position of the middle digit in the left front paw for single animal in each frame ^19^. The periodogram function of MATLAB was utilized along with in-house code to find the frequency of movement of the paw.

### Spatiotemporal gait analysis

Given that the mutant mice performed very poorly in a complex coordination task like the Rotarod, we chose to examine any abnormalities in gait using a simplified task associated with a lower level of stress. We therefore performed a quantitative spatiotemporal gait analysis of 3-month-old mice walking across an arena with a system involving high-speed videography ^20^. This approach increased hardware sensitivity with high-speed videography (250 frames per second), reduced the contribution of animal stress to gait measurements by avoiding a treadmill, and incorporated a second angle of view offering estimation of both spatial and temporal parameters. We manually digitized two-time variables (toe-off and foot-strike events) for each paw, and location coordinates (x,y) for each step. Animals underwent gait testing as described earlier by Dr. Kyle D. Allen’s group ^21^. The animals were acclimatized to the testing room for 1 hour and allowed to explore the gait arena without any external stimuli until 3 trials each were collected where the animal walked with an approximate constant velocity. High speed videography was utilized to collect walking videos at 250 frames per second (RedLake, M3). The collected videos were hand digitized by an observer using a modified version of DLT data viewer in MATLAB ^19^. Digitized data was then processed using MS Excel to calculate velocity, stride length, stance times, step widths along with spatial and temporal gait symmetry using methods described earlier ^20^. Velocity was included as a covariate for calculations of stride length, step width and percentage stance time analyses as per previous studies ^20^, and calculated residuals were utilized for statistical analyses.

### Tissue collection and sectioning

Mice were perfused with cold 4% paraformaldehyde using a Watson-Marlow 120S pump, and the cerebella were subsequently collected with post-fixation for 6 hours at 4°C. The cerebella were then subjected to consecutive sucrose treatments (15% and 30%) after which they were embedded in the Scigen Tissue-Plus™ OCT. 30um sections were cut using a cryostat starting from the midline and used for staining steps.

### Immunofluorescence staining, Cresyl Violet staining, and imaging

Cryostat sections were thawed in a humid chamber at room temperature and washed 2 x 3 min in 1x phosphate buffer saline (PBS). For immunofluorescence staining, sections were washed and rehydrated three times in 1x PBS. Permeabilization was performed with 0.25% PBS-Tween (PBS-T) and blocked with 3% serum in PBS-T for 1 hour. The sections were then subjected to primary antibodies: anti-calbindin (Abcam Cat# ab11426, 1:500) overnight at 4°C, followed by 3 x 5 min PBS washes and secondary antibody (Alexa 488 or Alexa 568, Thermo Fisher) treatment for 1 hour at room temperature. Coverslips were placed using the Vectashield mounting medium (Vector Laboratories). For Cresyl violet staining, the thawed and washed slides were then incubated in ddH20 and dipped in Cresyl violet solution for 20 min after which they were subjected to ethanol and xylene dehydration. Zeiss LSM 700 microscope was utilized to document the results and post-processed for summation of stacks using Zeiss Zen Lite and ImageJ.

### Electrophysiology

Sagittal cerebellar slices (300-μm thickness) were prepared from 1-month-old mice (wild-type: 4-5 slices per mouse, n=4, and R424H: 4-5 slices per mouse, n=4). The slices were then incubated for at least 1 hour in a preincubation chamber (Warner Instruments) at room temperature in conventional artificial CSF containing (in mM): 125 NaCl, 3 KCL, 2 CaCl2, 1 MgCl2, 1.25 NaH2PO4, 26 NaHCO3 and 10 glucoses continuously saturated with 95% O2 and 5% CO2. Cell-attached recordings were acquired with a MultiClamp 700B Microelectrode Amplifier in voltage-clamp mode at 33–34°C, by visually identifying Purkinje cell somata in anterior lobule III using Zeiss Axio Imager 2 Microscope (RRID:SCR_018876). The resistance of the electrode was 4 to 5 MΩ when filled with internal solution (in mM: 140 potassium gluconate, 5 KCl, 10 HEPES, 0.2 EGTA, 2 MgCl2, 4 MgATP, 0.3 Na2GTP and 10 Na2-phosphocreatine, pH 7.3 with KOH). The holding potential was 0 mV. Signals were filtered at 2 kHz and sampled at 10 kHz. On-line data acquisition and off-line data analysis were performed using Clampfit.

### Statistical analysis

All data are presented as mean ± standard error of mean. For spatiotemporal gait analysis, residual changes were calculated for stride length, stance time and step widths with velocity as a covariate and wild-type mice as controls in MS Excel. Resulting spatiotemporal gait data and other behavioral analysis data was compared using the Mann Whitney U-test or unpaired t-test with Welch’s correction in GraphPad Prism. For comparisons involving multiple time-points and genotypes, one-way ANOVA was performed in GraphPad Prism. Statistical significance was defined as p < 0.05 for all analyses.

## Results

### Generation of BAC-R424H mice

The cloning and recombineering strategy were summarized in Fig.1A. To confirm the mutation and generation of R424H mice, our genotyping data showed a clear DNA product (631 bp) in affected mice (Fig 1.B). Furthermore, quantitative PCR of cerebellum of one month old BAC-R424H mice confirmed the significantly decreasing mRNA expression levels of *Kcnc3* and calbindin (Fig 1.C).

**Figure 1.**
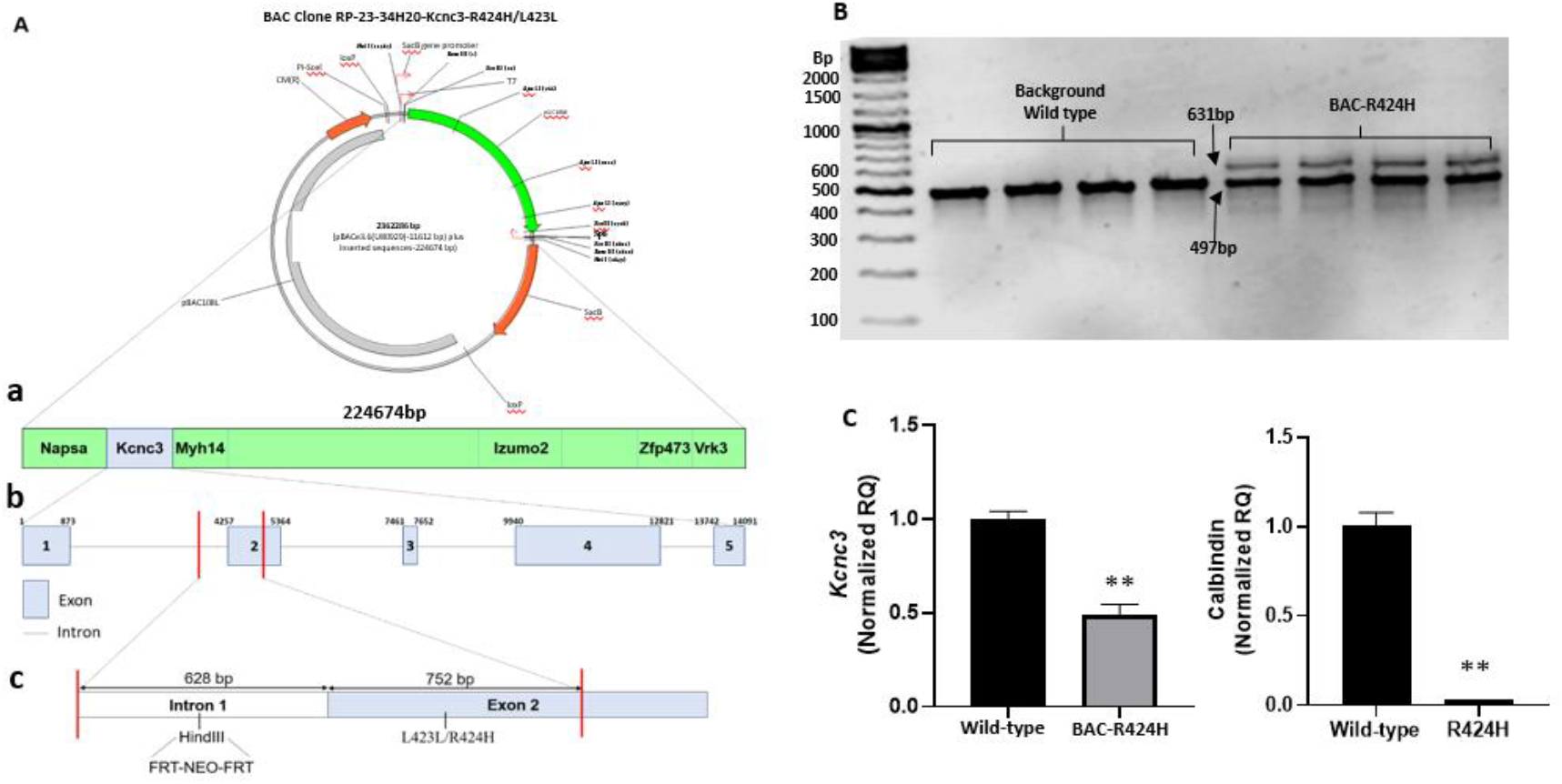
Generation of BAC-R424H mice. (A) A plasmid restriction map of BAC clone containing an inserted 224 674 bp mouse genomic fragment encoding the *Kcnc3* locus. (a) The order of the genes encoded in RP-23-34H20 BAC in an approximate scale. (b) *Kcnc3* intron exon structure. (c) Intron1/Exon2 fragment highlighting the location of the NEO cassette insertion in intron 1 and the L423L/R424H mutagenesis site. (B) PCR genotyping confirmed BAC-R424H mice, carriers displayed 631bp band. (C) Data of qPCR for *Kcnc*3 and Calbindin from wild-type mice and BAC-R424H mice. Note: ** p <0.01, n=5 per group.

### Significant neurological function impairment in BAC-R424H mice

BAC-R424H mice showed abnormal behavioral symptoms including abnormal power spike in frequency of tremor and aberrant gait, observed as early as at 2 weeks of age and persisting through adulthood. The relative power of frequencies of fore-paw tremor of 3-month-old mice were estimated using high-speed videos and power spectral analyses. The 3-month-old BAC-R424H mice showed high frequency tremor with a powerful spike at 46.35 ± 4Hz, as compared to age-matched wild-type mice that did not exhibit this frequency of limb movement. (Fig 2.A, B, and Suppl Video 1).

**Figure 2.**
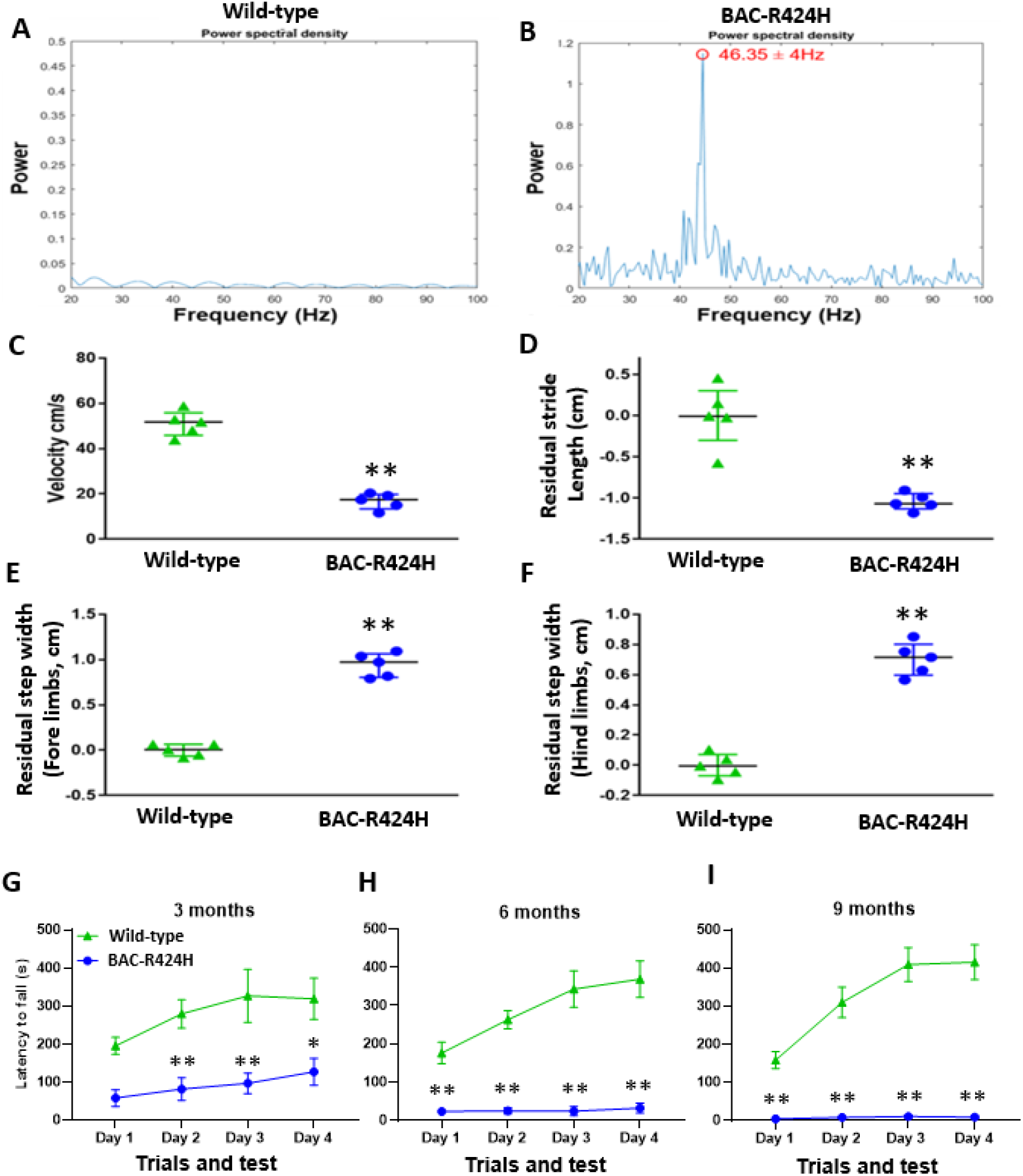
Neurological function impairment of BAC-R424H mice. (A-B) Tremor analysis in wild-type control mice and BAC-R424H mice: (A-B) Periodograms describe the relative power of frequencies present in data collected from fore paw tremors of wild-type (n=5, trials = 2) and BAC-R424H mice (n=5, trials = 2). The powerful spike of frequencies is difference between the wild-type mice (A, normal power spectrum), and the BAC-R424H mice (B, an abnormal power spike in frequency at 46.35±4Hz). (C-F) Spatiotemporal gait analysis in wild-type control mice and BAC-R424H mice: (C) Velocities of locomotion, (D) Residual stride length (cm), (E) Residual step widths (cm) of fore, and (F) Residual step widths (cm) of hind limbs. (G-I) Rotarod testing in wild-type control mice and BAC-R424H mice: Coordination on a rotating rod was evaluated for mice at ages of 3 months (G, n = 5 per group), 6 months (H, n = 6 per group) and 9 months (I, n = 7 per group). Note: One-way ANOVA Turkey multiple comparison test was used to analyze the statistical differences about the latency to fall between wild-type mice and BAC-R424H mice. ** p <0.01, * p <0.05.

Analysis of spatiotemporal gait patterns indicated that the BAC-R424H mice displayed aberrations. The 3-month-old BAC-R424H mice showed a significantly lower velocity and notably different residual stride lengths compared to those of wild-type mice (Fig 2.C, D). Major differences in step widths of both fore and hind limbs were observed in mutant mice, compared to wild type (Fig 2.E, F, and Suppl Video 2).

We were unable to perform the gait analysis on older mice (8 months) due to their inability to ambulate through the arena. In fact, most of them displayed an inability to balance their weight leading to multiple falls in the duration of the task. In summary, the BAC-R424H mice at 3 months of age displayed significantly slower ambulation, with shorter strides and wider steps in both fore and hind limbs, along with a lack of sustained coordination in gait cycles.

Accelerating Rotarod was used to test balance, grip strength and motor coordination of mice. At 3 months, the BAC-R424H mice showed marked impairment in their ability to stay on the Rotarod in comparison with wild-type mice on the day of testing (Fig 3.G). By 6 months, the BAC-R424H mice showed a significant deterioration in performance, implying a progression in neurologic functional impairment (Fig 3.H). At 9 months, the BAC-R424H mice were barely able to stand on the rod, exhibiting highly significant differences in motor coordination relative to the wild-type mice (Fig 3.I). Because of the severity of disability, no BAC-R424H mice can survive more than twelve months.

**Figure 3.**
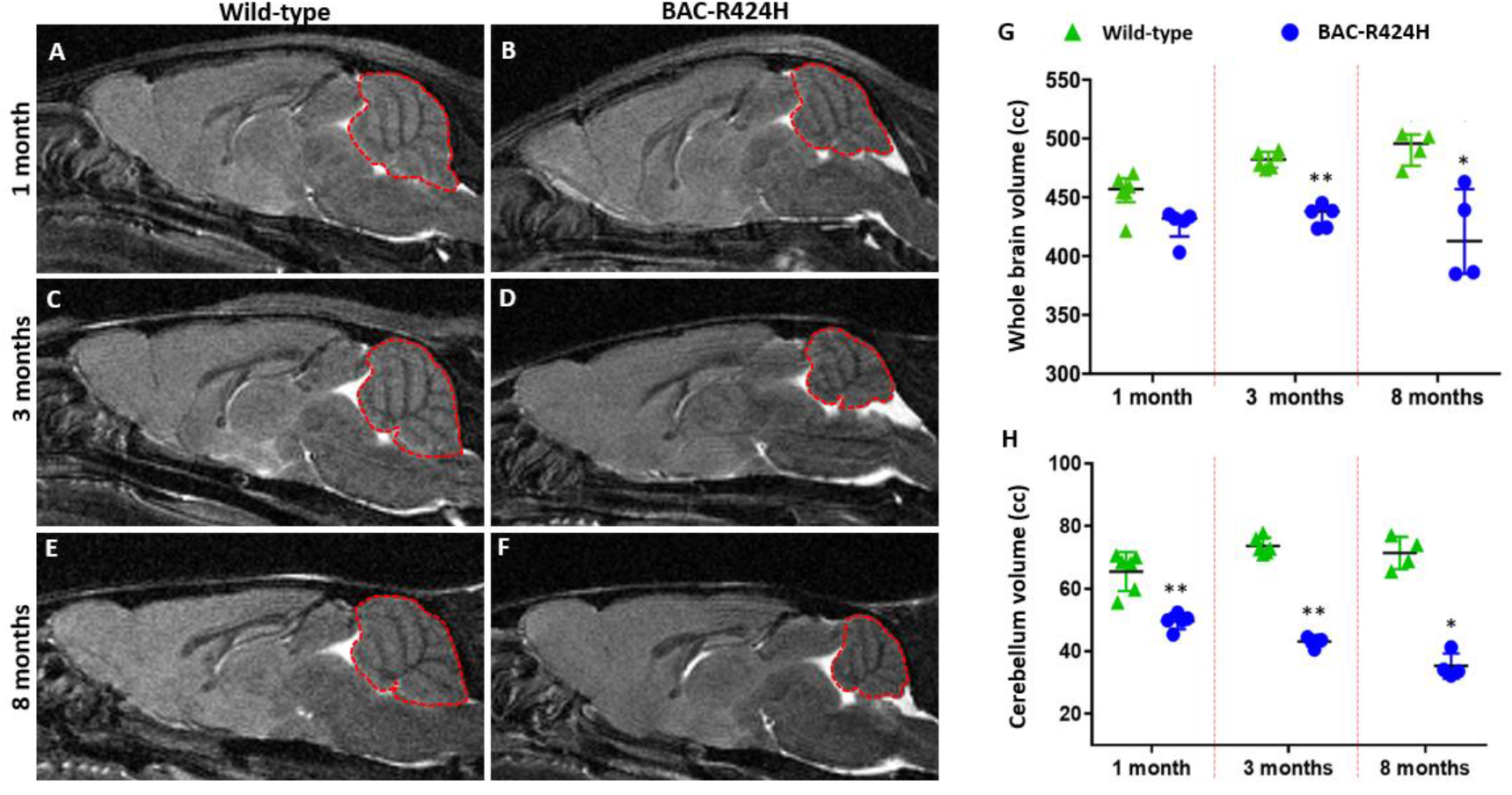
Cerebellar aggressive atrophy in BAC-R424H mice. Representative 7T MRI images of wild-type and BAC-R424H mice at ages of 1 month (A, B), 3 months (C, D), and 8 months (E, F). (G) The volumes of whole mice brains at 1 month (n = 5-6), 3 months (n = 5-6) and 8 months (n = 4). (H) The volumes of mice cerebellum. Different mice were used, and data was collected at every time point. Note: Graphs represent median, interquartile range from Mann-Whitney statistical analysis for each group. ** p <0.01, * p <0.05.

### Cerebellar hypoplasia and progressive atrophy in BAC-R424H mice

Patients with SCA13 R423H show marked cerebellar hypoplasia with MRI at ages as young as 10 months ^10^. To determine if the BAC-R424H mice also demonstrated developmental hypoplasia, we utilized 7T MRI to estimate the volumes of whole brains and cerebella. MR imaging showed the cerebella of BAC-R424H mice were significantly smaller than those of wild-type mice (Fig 3. A-F). The quantitative analysis of MRI data indicated that our BAC-R424H mice showed 6.6-14.3% decrease in whole brain volume when compared to controls at 1, 3 and 8 months of age (Fig 3. G). The cerebellar volumes of BAC-R424H mice were significantly smaller, exhibiting about a 28% decrease in volume at one month, 40% at three months, and 50% at eight months compared to the wild-type mice (Fig 3.H). The data indicated a progression of atrophy in addition to the developmental hypoplasia seen at one month.

### Neurodegeneration in BAC-R424H mice

We conducted a systematic assessment of the cerebellar molecular and granule cell layers to further investigate the etiology of the observed cerebellar atrophy. Molecular layer thickness was measured in Nissl-stained sections of mice cerebella (Fig 4.A, B). Our data show that at age as early as one month, the molecular layer thickness in cerebella of BAC-R424H mice was reduced significantly compared to wild-type mice (∼30% reduction), with progressive and significant loss in thickness at three months (∼50% reduction) and eight months (∼55% reduction) (Fig 4.C). A similar assessment of the granule cell layer however demonstrated no significant changes in the thickness of this layer in both mice groups.

**Figure 4.**
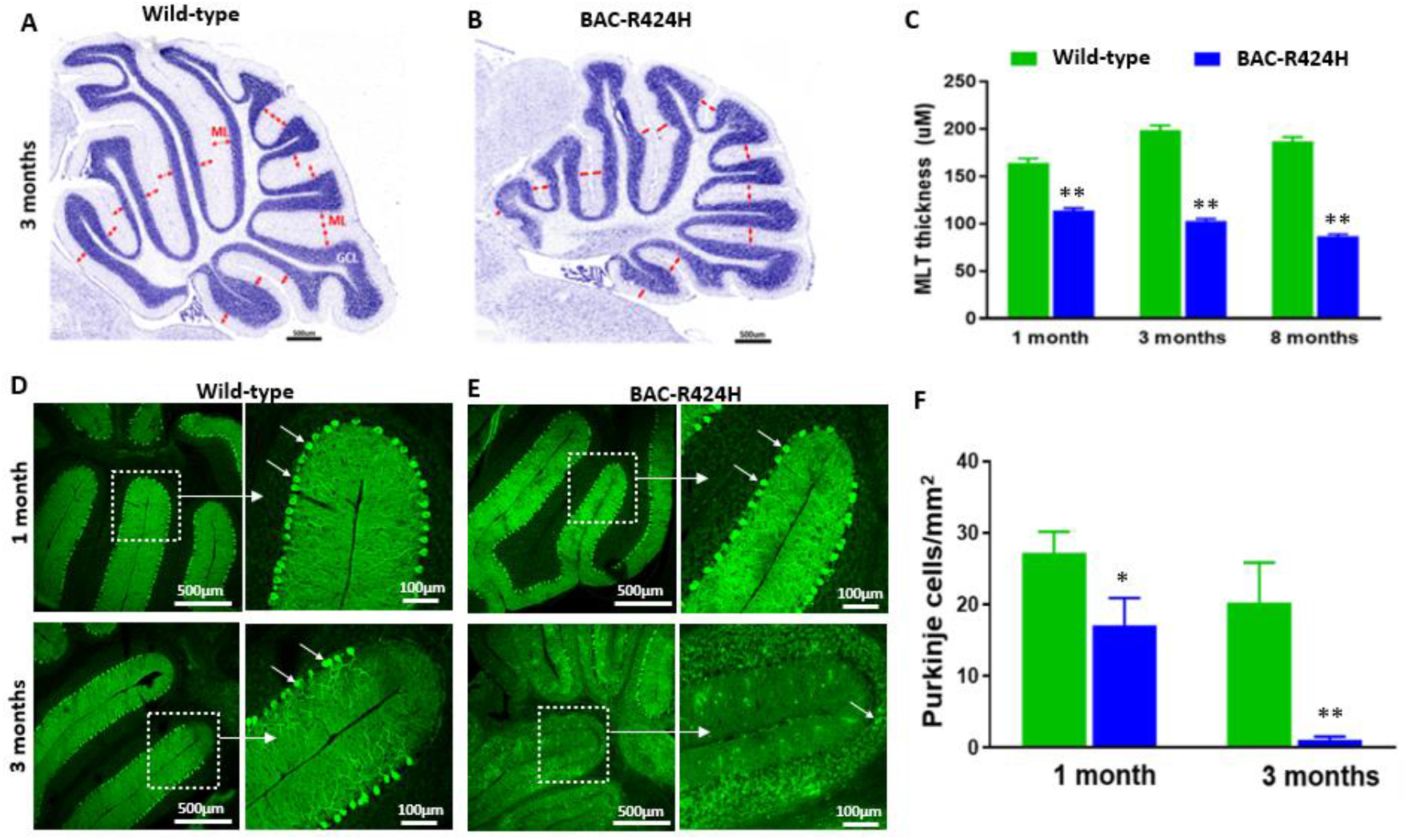
Cerebellar hypoplasia in BAC-R424H mice. (A-C) Representative images of Cresyl violet staining and analysis of molecular layer (ML) thickness in mice cerebellum. Two measurements were made for each lobe of the cerebellum, as indicated by the red arrows (A, B), and analysis of molecular layer thickness (C) of mice at 1 month (n = 4 per group), 3 months (n = 4 per group), and 8 months (n = 3 per group). (D-F) Representative images of Calbindin positive Purkinje cells in mice cerebellum at 1 month and 3 months. Calbindin positive Purkinje cells (white arrow) in cerebellum of wild-type mice (D) and BAC-R424H mice (E). Quantitative analysis of Calbindin positive Purkinje cells/mm^2^ in the cerebellum of all mice (n = 4 per group, 3 slices each mouse). Note: Data was presented as mean ± SEM and analyzed using one-way ANOVA and Tukey’s HSD post-hoc analysis. ** p <0.01, * p <0.05.

Purkinje cell loss is thought to be a major pathological characteristic in cerebella of patients with SCA13 ^22–24^. To corroborate these observations, we quantified the loss of Calbindin positive Purkinje cells in BAC-R424H mice (Fig 4.D, E). Our data revealed that the number of Purkinje cells in cerebella of BAC-R424H mice was significantly lower at one month (p<0.05) and three months (p<0.01), compared to wild-type mice (Fig 4.F). At eight months of age, BAC-R424H mice showed nearly complete loss of Purkinje cells, making it almost impossible to perform any meaningful and accurate counting analysis to compare wild-type mice and BAC-R424H mice at this time point.

### Abnormal spontaneous firing of Purkinje cells of BAC-R424H mice

Finally, we also analyzed the spontaneous firing properties of Purkinje cells based on the electrophysiological recordings in sagittal cerebellar slices from wild-type mice and BAC-R424H mice at the age of 1 month (Fig 5. A, B), prior to the previously observed extensive cell loss. The firing frequency of Purkinje cells in BAC-R424H mice (Fig 5. C, ∼30Hz) was significantly reduced when compared to that of the wild-type mice (Fig 5. C, ∼75Hz).

**Figure 5.**
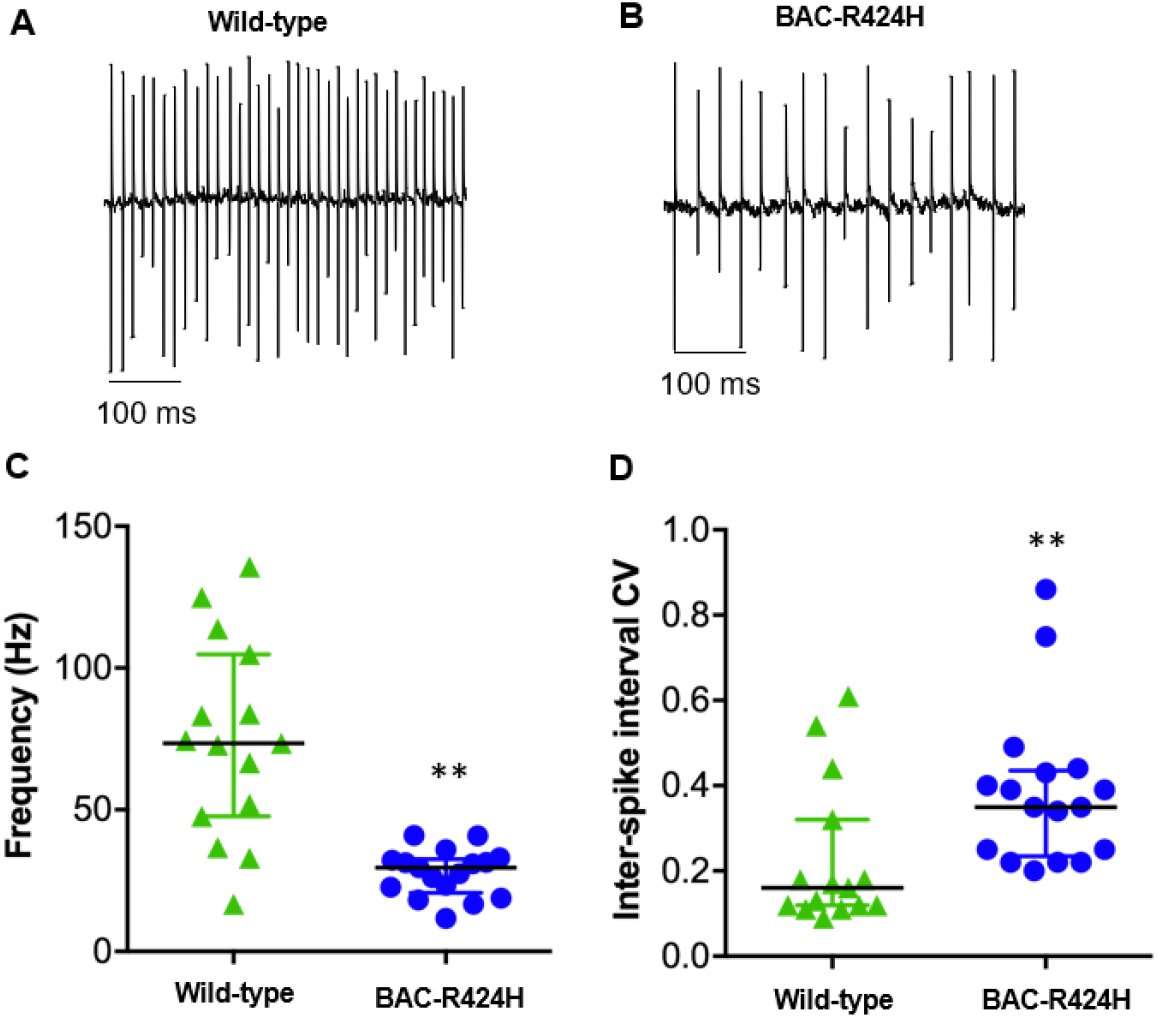
Abnormal spontaneous firing of Purkinje cells of BAC-R424H mice. Sagittal cerebellar slices were prepared from 1-month-old mice. Cell-attached recordings were acquired with a Multiclamp 700B amplifier in voltage-clamp mode by visually identifying Purkinje cell somata in anterior lobule III using an upright microscope (Zeiss). On-line data acquisition and off-line data analysis were performed using Clampfit. (A) Representative trace from wild-type mice cerebellar slices, (B) Representative trace from BAC-R424H mice cerebellar slices. (C) Spontaneous firing frequency of Purkinje cells, (D) Co-efficient of variation (CV) of the Inter-spike intervals (ISI) of Purkinje cells. Note: Data was presented as mean ± SEM and analyzed using Mann Whitney U-test. Note: wild-type, 4-5 slices per mouse, n=4, and BAC-R424H: 4-5 slices per mouse, n=4. ** p <0.01.

Correspondingly, the coefficient of variation of inter-spike interval was significantly increased, suggesting a change of firing pattern in BAC-R424H mice (Fig 5. D).

## Discussion

We report here the generation and characterization of a novel mouse model of the R423H allelic form of SCA13. This new BAC-R424H model recapitulates similar clinical SCA13 characteristics including both behavioral and neuropathological changes. Our BAC-R424H mice developed early onset clinical symptoms with obvious aberrant gait, tremor, and significant cerebellar hypoplasia and progressive atrophy.

One of the significant findings was the presence of a high-frequency tremor in the mutant mice, observable as early as within the first two weeks of life. In humans, tremor has a wide number of classifications including essential, dystonic, intention, and Parkinsonian. Essential tremor frequencies in humans range from 5-8Hz ^25^ whereas Parkinsonian tremor frequencies are around 4-7Hz ^26^. In our observations of the BAC-R424H mice, we identified that the strongest tremor was appendicular, especially in the upper extremities. Therefore, we evaluated fore-paw tremor frequencies in 3-month-old mice and demonstrated an average frequency of ∼46Hz. Tremor is also reported in this allelic form of SCA13 in patients, in fact it can be an impediment to motor activity requiring direct address. Gait ataxia is also a fundamental feature of SCA13 ^20^. Based on spatiotemporal gait analysis, we have been able to demonstrate that the 3-month BAC-R424H mice walked significantly slower than the wild-type cohort, with shorter strides and wider steps in both fore and hind limbs. These mice also lack sustained coordination in gait cycles. By 8 months of age, most mice display such an inability to balance their weight that gait analysis is impossible to accurately perform due to multiple falls and unsuccessful navigation through the arena during the task.

Our mouse model provides a first look at the pathology associated with the *Kcnc3* R423H mutation in the cerebellum at three different stages (1, 3, and 8 months) of development and maturation in mammals. An analysis of cerebellar histopathology shows an early change in Purkinje cell function with alterations in morphology and their significant loss as early as one month postnatal. There was progressive loss of Purkinje cells and decrease of molecular layer thickness, leading to notable cerebellar atrophy at 3- and 8-month-old BAC-R424H mice. Interestingly, immunohistochemistry staining of activated microglia clearly showed that most of activated microglia located around the Purkinje cells layer and significant astrocytes activation in the transgenic mice. This data indicated that the progressive Purkinje cells loss was accompanied by increased activation of microglia and astrocytes possibly associated with an ongoing inflammatory response.

Additionally, an abnormal increase in the expression of calbindin, the calcium binding protein, in the Bergmann glia of the cerebellum was observed. Previous studies have indicated an increase in the expression of calbindin in glia of other brain regions after ischemia or because of neurovirulence ^27, 28^. One of several functions of glia is their role in adapting to extracellular stresses including ionic imbalance, especially with potassium ion spatial buffering ^29^. Future studies analyzing the potassium and calcium ionic imbalances in the cerebellum, specifically in the Purkinje neurons and the Bergmann glia may reveal the order of events in SCA-associated cerebellar atrophy. Clues to neurodegeneration may be applicable to other brain regions and pathologies. Though there are several other genes contained within our BAC, only Myh14 is expressed in the CNS. The proteins of Napsa and Vrk3 are not detectable in the brain. We followed the MIQE protocol for RT qPCR and although apparently amplified, both Napsa and Vrk3 were below the level considered background. While in the transgenic mice with the inserted BAC construct including these two genes, we observed the two-fold increased mRNA levels after normalization. The fact that its expression persists through the lifetime of the BAC-R424H mice supports the assertion that the observed pathology is attributable to the mutant *Kcnc3* allele.

The frequency of intrinsic firing activity of Purkinje cells in our BAC-R424H model mice shows a significant reduction, which was also observed in mice models of other SCAs ^30–36^. *Kcnc3* R424H expression through lentiviral transduction revealed altered physiology and pathology of the cerebellar Purkinje cells with a degeneration in the viability in mouse primary cultured Purkinje cells ^37^. Electrophysiology recordings of these transduced cells showed severe suppression of current amplitude, with a broadening of action potentials and an increase in intracellular Ca2+ levels. Importantly, Purkinje cells expressing this R424H isoform showed morphological changes with smaller soma and shorter, less elaborate dendrites, in addition to accelerated cell death ^37^. The loss of Kcnc3 in Purkinje cells resulted in broadening of action potentials, with a reduction in frequency of spikelets ^15, 38^. Electrophysiological analysis of Xenopus oocytes expressing *Kcnc3* R423H deemed the channel a null with no K+ current conductance ^8^. The absence of current may be explained by biochemical data in mammalian cell culture whereby mutant channels are intracellularly retained in Golgi/endosomal vesicles, with limited plasma membrane localization ^9, 10^. Furthermore, upon co-expression of differing ratios of R423H and wild type channels, we observed a dominant negative effect with intracellular sequestration of both channels ^10^. Studies in Xenopus oocytes showed differences in kinetic behavior of predicted heteromeric R423H wild type channels, with slower inactivation as compared to wild type channels ^39^.

In this study, we observed significantly reduced expression of Kcnc3 in the transgenic mice compared to wild type controls. Kcnc3 gene encodes the Kv3.3 voltage dependent potassium channel, the R424H mutation in Kcnc3 gene impaired the function of potassium channel and leaded to the dysfunction of Purkinje cells including abnormal spontaneous firing, and eventually its degeneration and loss. It suggests that the R424H mutation may have a dominant effect on the expression of endogenous Kcnc3, but the fully pathological molecular mechanism induced by R424H mutation remains unclear. Our previous *in vivo* Drosophila model of SCA13 expressing *KCNC*3 R423H also showed early-onset neurodegeneration in photoreceptors, manifesting as a maldeveloped eye. Interestingly, over-expression of the developmentally important epidermal growth factor receptor rescued the observed eye phenotype ^10^. However, the molecular mechanisms that lead to these deficits in this neurodegenerative allelic isoform model are not well understood. Future experiments involving transcriptomics, proteomics, and histopathological approaches in the early developmental stages of BAC-R424H mice may help us understand the developmental changes leading to cerebellar dysfunction and resultant neurodegeneration in the cerebellum and other cortical areas with increased expression of KCNC3.

In summary, we have developed a mouse SCA13 model of the causative, neurodevelopmental *Kcnc3* mutation R424H that uniquely displays both behavioral and pathological phenotypes. These transgenic mice exhibit the pathological and behavioral features of the homologous R423H mutation in human patients. Decrease in the cerebellar volume of BAC-R424H mice with concomitant loss of Purkinje cells and decline of molecular layer thickness from 3 to 8 months of age offers a venue for studying the neurodevelopmental and neurodegenerative features of this allelic form of SCA13. This transgenic model mimicking both neuropathological and behavioral changes manifested in SCA13-affected patients will provide a unique opportunity to understand the developmental role of voltage-gated potassium channels in cerebellar morphogenesis as well as insights into neurodegeneration seen in the cerebellum and additional brain regions. It is a robust model for investigating neurodevelopmental and degenerative processes *in vivo* from the molecular to the behavioral level and for evaluating potential therapeutic compounds.

## Supplementary materials (Attached)

## Acknowledgements

We appreciated the help from University of Michigan’s Biomedical Research Core Facility. They were the ones that verified the BAC sequence, grew and purified the BAC DNA, injected into fertilized oocytes, and identified and transferred the positive lines to us.

## Declarations

### Ethics approval

All experiments involving animals were performed following the protocols in accordance with the Revised Guide for the Care and Use of Laboratory Animals that was approved by the Institutional Animal Care and Use Committee (IACUC) of the Barrow Neurological Institute.

### Availability of data and material

The data that support the findings of this study are available from the corresponding author upon reasonable request.

### Authors’ contributions

Conceptualization: Michael F. Waters; Methodology: Jerelyn A. Nick, Swati Khare, Junxiang Yin, Heidi E. Kloefkorn, Ming Gao; Formal analysis and investigation: Junxiang Yin, Jerelyn A. Nick, Swati Khare, Heidi E. Kloefkorn, Ming Gao, Michael Wu, Jennifer White, Michael F. Waters; Writing - original draft preparation: Junxiang Yin, Swati Khare, Jerelyn A. Nick, Michael Wu, Jennifer White, Michael F. Waters; Writing - review and editing: Junxiang Yin, Swati Khare, Jerelyn A. Nick, Michael Wu, Jennifer White; James L. Resnick; Kyle D. Allen, Harry S. Nick, Michael F. Waters; Funding acquisition: Michael F. Waters. All authors read and approved the final manuscript.

